# WEVar: a novel statistical learning framework for predicting noncoding regulatory variants

**DOI:** 10.1101/2020.11.16.385633

**Authors:** Ye Wang, Yuchao Jiang, Bing Yao, Kun Huang, Yunlong Liu, Yue Wang, Xiao Qin, Andrew J. Saykin, Li Chen

**Affiliations:** Department of Medicine, Indiana University School of Medicine, Indianapolis, IN, 46202; Center for Computational Biology and Bioinformatics, Indiana University School of Medicine, Indianapolis, IN, 46202; Department of Computer Science and Software Engineering, Auburn University, Auburn, AL, 36849; Department of Biostatistics, Gillings School of Global Public Health, University of North Carolina, Chapel Hill, NC 27599; Department of Genetics, School of Medicine, University of North Carolina at Chapel Hill, Chapel Hill, NC 27599; Department of Human Genetics, Emory University, Atlanta, GA 30322; Department of Medical and Molecular Genetics, Indiana University School of Medicine, Indianapolis, IN, 46202; Department of Radiology and Imaging Sciences, Indiana University School of Medicine, Indianapolis, IN, 46202

**Keywords:** noncoding variants, prioritization, functional score

## Abstract

Understanding the functional consequence of noncoding variants is of great interest. Though genome-wide association studies (GWAS) or quantitative trait locus (QTL) analyses have identified variants associated with traits or molecular phenotypes, most of them are located in the noncoding regions, making the identification of causal variants a particular challenge. Existing computational approaches developed for for prioritizing non-coding variants produce inconsistent and even conflicting results. To address these challenges, we propose a novel statistical learning framework, which directly integrates the precomputed functional scores from representative scoring methods. It will maximize the usage of integrated methods by automatically learning the relative contribution of each method and produce an ensemble score as the final prediction. The framework consists of two modes. The first “context-free” mode is trained using curated causal regulatory variants from a wide range of context and is applicable to predict noncoding variants of unknown and diverse context. The second “context-dependent” mode further improves the prediction when the training and testing variants are from the same context. By evaluating the framework via both simulation and empirical studies, we demonstrate that it outperforms integrated scoring methods and the ensemble score successfully prioritizes experimentally validated regulatory variants in multiple risk loci.

## Introduction

In the past decade, genome-wide association studies (GWAS) have been widely used to identify tens of thousands of genome-wide significant tag SNPs associated with complex traits. However, tag SNPs may not be causal as the association is possibly mediated by a causal SNP associated with both the tag SNP and the trait. Nevertheless, it is difficult to determine the underlying causal variants due to complex patterns of LD among SNPs. Moreover, quantitative trait locus (QTL) analyses have successfully identified variants associated with molecular phenotypes i.e. gene expression, DNA methylation, chromatin accessibility [1, 2, 3, 4, 5]. These molecular QTL studies enable the understanding of molecular basis of GWAS SNPs via colocalization. However, the high sequencing cost leads to QTL studies with modest sample sizes, limiting the power to uncover QTLs with small effects. Therefore, identification of these functional noncoding variants that have direct functional consequence on complex traits and molecular phenotypes remains challenging in human genetics research.

Several studies suggest that functional noncoding variants are believed to disrupt the normal regulatory activity in promoter and enhancer regions in order to impact the downstream gene expression in a tissue or cell type specific manner and thus result in the onset of disease such as the prevalence of TERT promoter mutations has been established in melanoma, gliomas and bladder cancer [6]; novel MYB-binding motifs, which are generated by somatic mutations in the intergenetic regions, creates a super-enhancer upstream of the TAL_1_ oncogene in a subset of T cell acute lymphoblastic leukaemia [7]. Moreover, more than 90% of GWAS identified SNPs are noncoding and are enriched in regulatory elements (REs). A recent exploratory study demonstrates that active chromatin marks (e.g. H_3_K_27ac_ and H_3_K_4me1_), and repressive chromatin marks (e.g. H_3_K_9me3_ and H_3_K_27me3_) show different regulatory activities between a risk variant rs3024505 associated with type 1 diabetes and a benign variant rs114490664 [8]. This example indicates that RE activity can be used to distinguish the causal and non-causal SNPs.

The rapid development of massively parallel sequencing technologies enables the generation of thousands of “multi-omics” data, which are publicly available at large national and international consortia such as the Encyclopedia of DNA Elements (ENCODE) [9], Roadmap Epigenomics [10] and International Hu-man Epigenome Consortium [11]. These multi-omics data measures genome-wide regulatory activities such as histone modifications (e.g. ChIP-seq), methylation (e.g. methylation array, whole-genome bisulfite sequencing), chromatin accessibility (e.g. DNase-seq, ATAC-seq) and chromatin interactions (e.g. Hi-C) across hundreds of tissues and cell types. Using standard sequencing data processing protocols such as peak-calling, tissue- or cell type-specific REs and RE activities can be detected. Variant annotations are further created by overlapping variants and REs where the variant fall in [8, 12, 13, 14]. These annotations have been widely used as predictive features to develop computational methods for predicting functional noncoding variants [15, 16, 17, 18, 19, 20, 21], which adopt different computational methodologies, use different training variants and utilize different variant annotations. Among these methods, supervised learning approaches, such as GWAVA [15], CADD [16], DANN [17], FATHMM-MKL [18]), LINSIGHT [19]), FunSeq2 [20], are trained using labelled non-causal variants and causal ones, either putative or experimentally validated, to predict the probability of a give variant for being causal. Different from the supervised learning methods, a common practice of unsupervised learning approaches such as Eigen [21] performs direct aggregation of multi-dimensional variant annotations into one single functional score, which measures the functional importance of the variant, without a training step. Importantly, for most of the existing methods, genome-wide precomputed functional scores for known variants from 1000 Genomes Project [22] or gnomAD [23] are publicly available. Without the need to retrain the model, users can obtain these scores efficiently by providing a list of variants identifiers or genomic coordinates and utilize these scores directly for post-GWAS study i.e. fine mapping analysis. Usually, a larger score indicates the variant could potentially be more functional and the variant with the highest score is prioritized in a risk locus with LD-linked variants.

Nevertheless, without strong prior knowledge, it is difficult to choose which scoring method in real application among multiple methods developed for the same purpose. It is even more challenging to make the choice considering prediction performance of existing scoring methods has been shown poor concordance on the state-of-the-art benchmark datasets [24]. There are two possible reasons to explain the poor consistency. First, these methods are trained using different training variants and variant annotations to predict functional noncoding variants from different context (i.e. disease, tissue or cell types), making one method trained using variants from one context have suboptimal prediction for variants from another context. Second, they adopt different algorithms tailored to specific scenarios, limiting the generality. For example, GWAVA is developed using pathogenic regulatory variants from The Human Gene Mutation Database (HGMD) [25] and is thus used to predict pathogenic regulatory variants; FunSeq2 is trained using recurrence cancer somatic variants and is therefore specifically designed to predict noncoding regulatory variants in cancer. Considering the above challenge, given a variant without prior knowledge about its context and functional consequence, an ensemble approach that combines the predictions of all these methods in a weighted scheme could offer a more powerful prediction than each method. The weight of each individual scoring method, which reflects their contributions in the prediction task, can be adaptively learnt in different context, which improves the generality and flexibility.

We hereby developed a statistical learning framework “WEVar” (<u>W</u>eighted <u>E</u>nsemble framework for predicting functional regulatory <u>V</u>ariants) by integrating representative scoring methods in a constrained optimization approach, where the precomputed functional scores of these methods are treated as predictive features with two constraints: i) the summation of weights of existing methods are required to be one; ii) a *L*_2_-norm is further imposed on the weights for smoothing the weight estimation. There are several advantages of WEVar. First, WEVar is developed directly on top of precomputed functional score, which is an optimally integrative metric that represents for thousands of multi-omics functional annotations used by each scoring method. Using these functional scores directly will decrease the number of predictive features dramatically and thus avoid the challenge of high-dimensional data in the model development, that is, the sample size of labelled causal variants is fewer than the number of variant annotations. Second, WEVar leverages individual scoring method by adaptively learning the contribution of each one, which will up-weight the methods fit more in the current context and down-weight the others, and thus optimizes the prediction performance. Last but most importantly, WEVar has two modes: “context-free” and “context-dependent”. Context-free WEVar is used to predict functional noncoding variants from unknown or heterogeneous context. Context-dependent WEVar can further improve the functional prediction when the variants come from the same context in both training and testing set. Using simulation and real data studies, we demonstrate both WEVar modes outperform each individual scoring method on the state-of-the-art benchmark datasets. Importantly, context-dependent WEVar can further improve the functional prediction. We also show that WEVar can successfully prioritize experimentally validated regulatory variants associated with different traits and located in different risk loci.

## Results

The overview of WEVar is shown in Figure 1. First, we will perform a simulation study to evaluate the accuracy of weight estimation by WEVar for all integrated scoring methods and investigate whether the prediction performance of WEVar is improved compared to individual scoring method. Second, we will evaluate the context-free functional prediction and context-dependent functional prediction on the state-of-the-art benchmark datasets respectively. Third, we will apply WEVar to prioritize experimentally validated causal regulatory variants in multiple risk loci associated with multiple traits.

**Figure 1.**
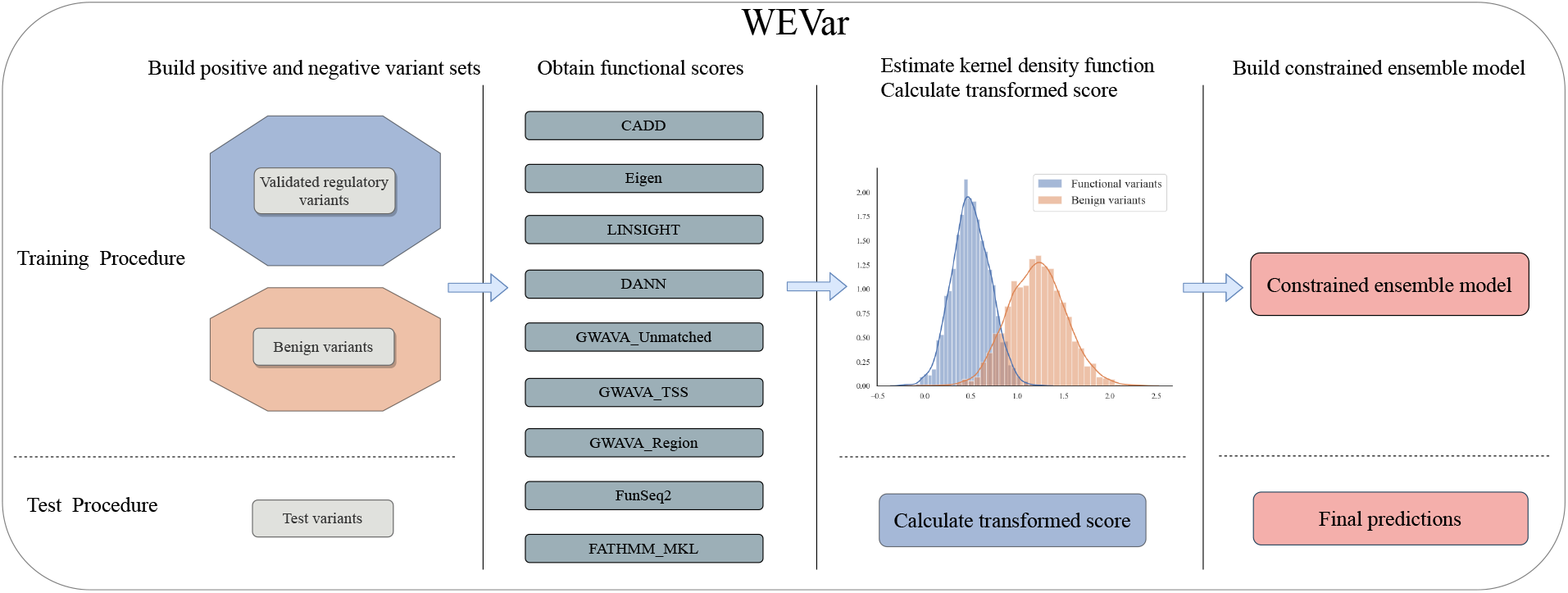
Overview of the WEVar. WEVar aims to predict functional noncoding variants, which has two modes: “context-free” and “context-dependent”. For “context-free” mode, the training variant set is chosen from a curated set of functional regulatory variants from diverse context to train a model for functional prediction of variants from unknown or heterogeneous context. For “Context-dependent” mode, the training variant set is selected from one specific context of interest (i.e. disease, tissue, cell type), to train a model for functional prediction of variants from the same context. In the training phase, WEVar compiles the training set with labelled functional and non-functional variants and annotate all variants with precomputed functional scores from representative scoring methods. For each method, the raw scores are transformed using kernel density function (KDE) for both functional and non-functional variant sets respectively. Using these transformed scores as predictive features, a constrained ensemble model is trained. In the testing phase, precomputed functional scores of testing variants are transformed based on the estimated KDE in the training phase and then serve as input features for trained ensemble model to predict the ensemble WEVar score.

### Evaluation of WEVar in a simulation study

#### Evaluation metrics

The performance of all scoring methods is evaluated using area under the receiver operating characteristics curve (AUROC), the area under the precision-recall curve (AUPR) and Pearson correlation between predicted and true labels (COR). AUROC and AUPR are metrics based on the ranks of the predicted scores. COR has the additional ability to measure how the predicted values are correlated with the true labels. Using different probability cutoffs, AUROC measures the trade-off between the true positive rate and false positive rate. AUPR compares the trade-off between the true positive rate and precision. AUROC is preferred for balanced class, whereas AUPR is more appropriate for imbalanced class. Since we have both balanced and unbalanced testing datasets, we present both metrics.

#### Simulating correlated functional scores and variant labels

We conduct a simulation study to evaluate whether WEVar can estimate contribution of each individual scoring method accurately and whether WEVar can improve prediction performance compared to each individual scoring method. Since the functional scores of different methods have an overall positive correlation (Figure 2A), we simulate functional scores of all scoring methods with consideration of the score correlation. Using the simulated scores, we generate a total 10, 000 variants with an equal size of functional and nonfunctional variants in the training set. Similarly, we independently generate an equal number of 10, 000 variants in the testing set for prediction evaluation. We then apply WEVar to retrain a model in the training set and predict WEVar scores in the testing set. Using WEVar scores and true labels in the testing set, we will calculate AUROC, AUPR and COR. We repeat the whole procedure 50 times and obtain the average of all evaluation metrics.

**Figure 2.**
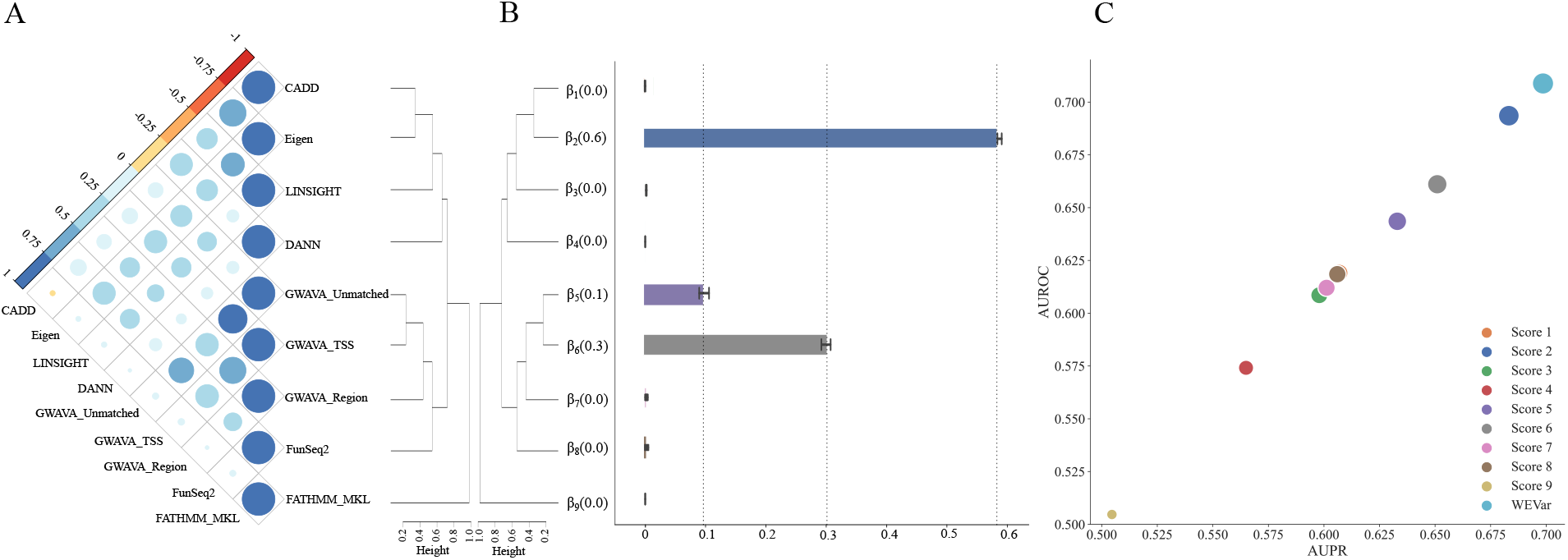
(A) Pairwise Pearson correlations between precomputed functional scores among scoring methods for the integrated causal regulatory variants collected from Li et al. [26]. (B) Average regression coefficient estimated by WEVar in the training phase in 50 simulations. (C) Average prediction performance by WEVar on the independent testing datasets. X axis presents AUPR; Y axis presents AUROC; the bubble size represents COR. AUPR, AUROC and COR are averaged in the testing phase in 50 simulations.

Specifically, using the integrated causal regulatory variant set collected from Li et al. [26], we calculate a *p* × *p* variance-covariance matrix *R* of precomputed functional scores among all integrated scoring methods, where *p* is the number of scoring methods. We cluster these methods based on Pearson correlation and find that these methods have different levels of disagreement, indicating that performance of these methods show poor concordance on the benchmark dataset (Figure 2A). Not surprisingly, GWAVA_Unmatched, GWAVA_Region and GWAVA_TSS are clustered together since they use the same positive training variant set. Surprisingly, FATHMM-MKL has the lowest correlation with all the other methods. Indeed, this observation highlights the rationale why a weighted ensemble strategy proposed by WEVar is essential to improve the prediction because it is able to upweight the scoring methods fit in current context while down-weight the unfit others. We further perform Cholesky decomposition on *R* as:

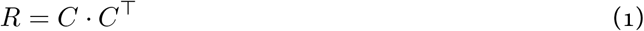

where *C* is a *p* × *p* lower triangular matrix with real positive diagonal entries. To maintain the correlations of simulated scores, we generate the correlated functional scores *X* as the product between *C*^T^ and random variable *d*, which is sampled from an independent normal distribution as:

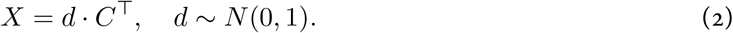

where *x_ij_* as the functional score of *ith* noncoding variant in *jth* scoring method. *η_i_*, which is the weighted average score of *ith* variant, can be generated as:

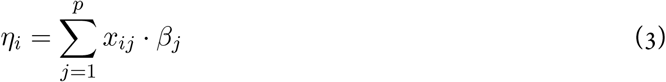

where *β_j_* is the weight associated with *j*th method. Without loss of generality, we manually assign 0.6 to *β*_2_, 0.3 to *β*_6_, 0.1 to *β*_5_, and 0 to the rest. We then perform inverse logit transformation to *η_i_* to obtain probability *π_i_*, based on which the binary label *y_i_* for *ith* variant is generated from a Bernoulli distribution as:

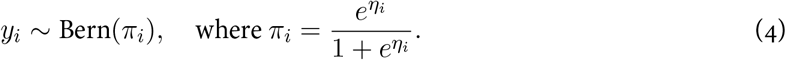

#### Results of the simulation study

In the simulation study, we will evaluate whether WEVar can truly discover the contributions of individual scoring method by comparing the estimated regression coefficients 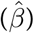 with the assigned true values (*β*). To fit a WEVar model, the optimal tuning parameter for *L*_2_-norm is selected using fivefold cross-validation (5-CV), where the whole training set is divided into five-folds, where four-folds is used to train the model and one-fold is used to obtain the evaluation metric i.e. AUROC. The optimal tuning parameter is chosen based on the average AUROC from 5-CV, and a final model is fitted using the whole training set with the optimal tuning parameter. To evaluate the performance of the final model an independent testing set, we use all evaluation metrics AUROC, AUPR and COR.

As a result, we find that the estimated weights are nearly unbiased to the underlying truths (Figure 2B), which suggests that WEVar can discover the contribution of each individual scoring method correctly when the functional scores of these methods are correlated. With accurate contribution estimation, WEVar can significantly improve the prediction performance in the independent testing (Figure 2C) by achieving the highest AUROC, AUPR and COR. Overall, the simulation results validate the benefit of exploiting different scoring methods in an integrative weighted scheme.

### Context-free functional prediction

#### Overview of context-free WEVar

We first introduce context-free WEVar, which is trained using integrated causal regulatory variants collected from Li et al. [26]. We call this WEVar mode “context-free” because these variants are not limited to a specific context but have a broad definition of functionality across a wide range of context. These variants are either experimentally validated or highly putative causal variants associated with different diseases, molecular phenotypes or clinical outcomes, which are located in different noncoding regions such as promoters, enhancers, _5_’UTRs and _3_’UTRs. The diverse context and widespread genomic locations of these variants make it potentially powerful to predict functional noncoding variants when the context is unknown or heterogeneous. To demonstrate the generality of context-free WEVar, we evaluate it on the independent benchmark datasets containing noncoding variants of different functionalities and from diverse context. We also remove any duplicated variants overlapped with training dataset from each independent testing dataset, which can prevent potential overfitting. To verify the effectiveness of the weight strategy, besides all scoring methods WEVar integrates, we also include “Unweighted average” as a comparison, which is the unweighted average of min-max normalized precomputed functional scores from the integrated methods. In the training phase of WEVar, tuning parameter for *L*_2_-norm is selected using 5-CV. For all methods, AURPC, AUCPR and COR are reported on each independent testing dataset.

#### Results of functional prediction between context-free WEVar and integrated scoring methods

We start to compare the prediction performance between WEVar and its integrated scoring methods on three datasets, which consist of putatively functional variants based on statistical association (Figure 3). The first dataset, which is produced by Maurano et al. [28] and processed by Li et al. [26], contains 8,592 significant allelic imbalanced SNPs of chromatin accessibility (FDR<0.1) as the positive set and 9,678 frequency-matched background SNPs around nearest transcription start sites of randomly selected genes as the negative set. We observe that WEVar obtains the largest AUROC, AUPR and COR (0.894, 0.852, 0.644) with substantial improvements over each individual scoring method (Table S1). Following WEVar, LIN-SIGHT, GWAVA_Unmatched and Unweighted average have an overall comparable performance. However, the COR of LINSIGHT is much lower (0.255) compared to GWAVA_Unmatched (0.535) and Unweighted average (0.559). Surprisingly, FATHMM_MKL also has the lowest COR (0.053). Moreover, CADD and DANN, which utilize the same training set, have comparable but poorest performance among all methods (CADD: 0.639, 0.610, 0.228; DANN: 0.634, 0.563, 0.236). Interestingly, the prediction performance of GWAVA_Unmatched, GWAVA_TSS and GWAVA_Region are discordant even if they use the same positive training set (GWAVA_Unmatched: 0.875, 0.823, 0.535; GWAVA_TSS: 0.840, 0.796 0.559; GWAVA_Region: 0.723, 0.691, 0.382).

**Figure 3.**
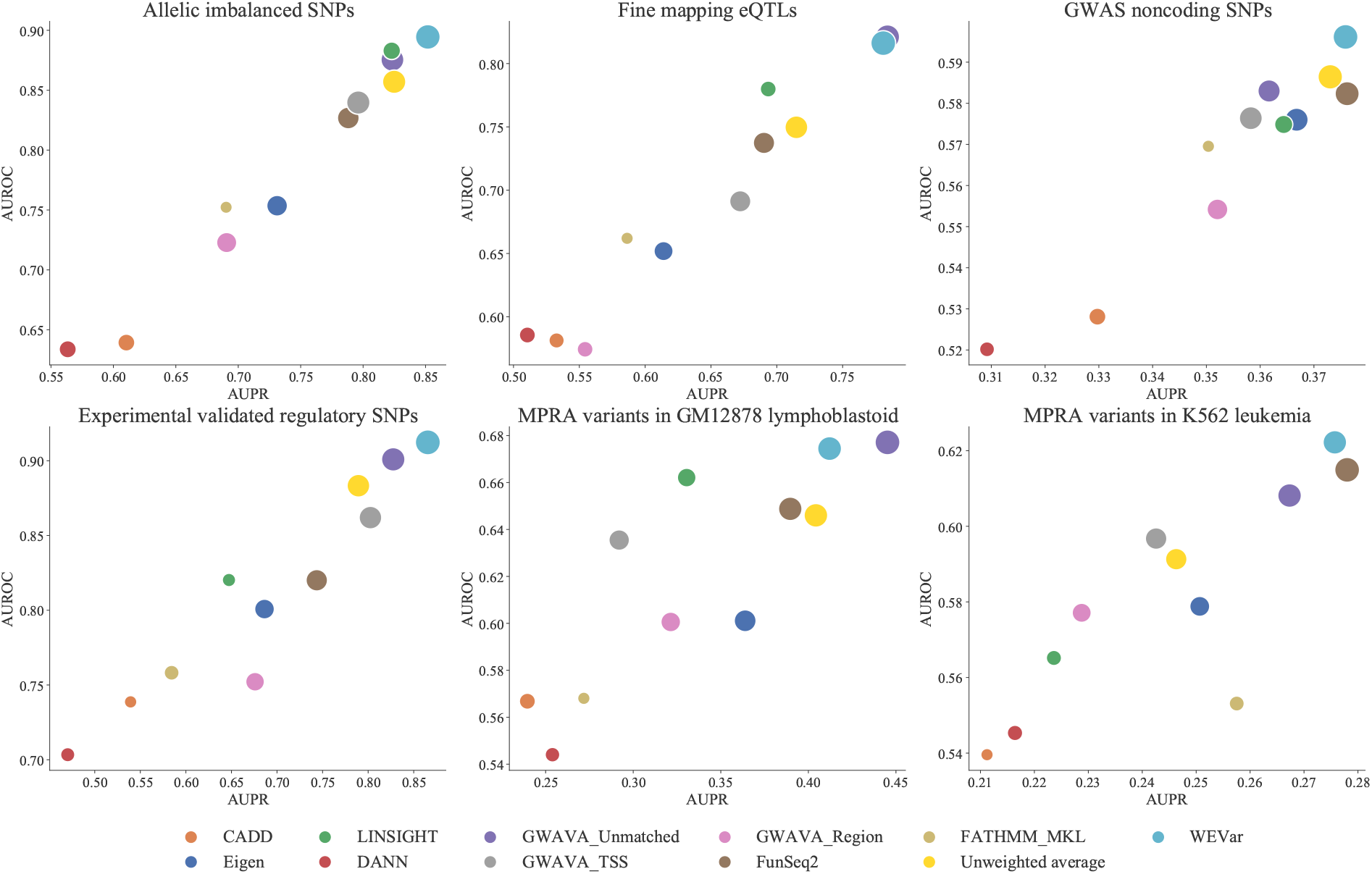
Evaluation of context-free WEVar and integrated scoring methods. Context-free WEVar is trained using the integrated functional regulatory variants collected by Li et al. [26], which include variants in HGMD, ClinVar, OregAnno and fine-mapping candidate causal SNPs for 39 immune and non-immune diseases with a total of 5,247 positive variants and 55,923 negative variants. Context-free WEVar is tested on the state-of-the-art benchmark datasets, which include i) Allelic imbalanced SNPs in chromatin accessibility with a total of 8,592 positive variants and 9,678 negative variants (Allelic imbalanced SNPs); ii) Uniformly processed fine-mapping eQTLs from 11 studies with a total of 31,118 positive variants and 36,540 negative variants (Fine mapping eQTLs); iii) GWAS noncoding SNPs with a total of 19,797 positive variants and twice number of negative variants (GWAS SNPs) [27]; iv) Manually curated experimentally validated regulatory SNPs with a total of 76 positive variants and 156 negative variants (Experimentally validated regulatory SNPs); v) MPRA validated variants in lymphoblastoid cells with a total of 693 positive variants and 2,772 negative variants (MPRA variants in GM12878 lymphoblastoid); vi) MPRA validated variants in erythrocytic leukemia cells with a total of 342 positive variants and 1,368 negative variants (MPRA variants in K562 leukemia). We further remove variants on sex chromosome or with missing precomputed scores. X axis presents AUPR; Y axis presents AUROC; the bubble size represents COR.

The second dataset consists of eQTLs in 11 studies across 7 tissues identified from Brown et al. [29] and processed by Li et al. [26]. The positive set consists of 31,118 significant eQTL SNPs (FDR<0.1) and the negative set contains 36,540 frequency-matched background SNPs around nearest TSS of randomly selected genes. We observe that WEVar has the largest COR and comparable AUROC and AUPR to GWAVA_Unmatched (WEVar: 0.816, 0.781, 0.509; GWAVA_Unmatched: 0.821, 0.781, 0.476) (Table S2). Moreover, both WEVar and GWAVA_Unmatched have clearly advantages over other scoring methods. For example, they improve nearly 0.04 AUROC and 0.09 AUPR over LINSIGHT, and 0.07 AUROC and 0.07 AUPR over Unweighted average. Particularly, there is substantial improvement of nearly 0.1 COR to Unweighted average and over 0.3 to LINSIGHT. Notably, the relative performance of GWAVA_Region drops dramatically and it has the lowest AUROC (0.574). FATHMM_MKL still has the lowest COR (0.047) followed by CADD and DANN (0.126, 0.1500).

The third dataset collects 19,797 GWAS significant noncoding SNPs from NHGRI-EBI GWAS Catalog [30] as positive set and twice number of variants in the negative set, which are randomly sampled from all noncoding variants in 1000 Genomes project with minor allele frequency (MAF) ≥ 5% [27]. The relative prediction performance of all methods are similar to the first dataset of allelic imbalanced SNPs. WEVar outperforms all scoring methods by obtaining the highest AUROC, AUPR, and COR. FATHMM_MKL have the lowest COR, while CADD and DANN have the lowest AUROC and AUPR (Table S3).

In addition to the three datasets comprised of putatively functional noncoding variants derived from association analyses, we compare the prediction performance between WEVar and all scoring methods on three datasets consisting of experimentally validated regulatory variants. The first dataset include 81 experimentally validated regulatory SNPs curated by Li et al. [26]. We find the trends of prediction performance for all methods still holds similarly to allelic imbalanced SNPs and GWAS significant noncoding SNPs, where WEVar obtains the largest AUROC, AUPR and COR (0.912, 0.865, 0.718) followed by GWAVA_Unmatched (0.901, 0.828, 0.649) and Unweighted average(0.883, 0.789, 0.617) (Table S4).

The other two datasets contain processed causal regulatory variants validated by MPRAs in two cell lines [31]. The first MPRA dataset includes 665 variants with genomic loci annotation in Ensembl database as positive set, which are selected out of 842 expression-modulating variants that show significantly differential allelic expression in GM12878 lymphoblastoid cells [32]. The negative set contains 2,772 control variants tested by MPRA but neither allele showed significant effects on expression (Bonferroni corrected pvalue>0.1). The second MPRA dataset consists of 339 positive variants that cause significant change of expression via targeted motif disruption in enhancers in K562 erythrocytic leukemia cells (pvalues<0.05) [33]. The negative set contains 1,359 control variants without causing significant change (pvalues>0.1). As a result, WEVar has comparable performance with top-performed GWAVA_Unmatched in predicting MPRA validated regulatory variants in GM12878 lymphoblastoid cells (WEVar: 0.674, 0.412, 0.286 vs GWAVA_Unmatched: 0.677, 0.445, 0.317) (Table S5). WEVar achieves largest AUROC and AUPR in predicting MPRAs validated regulatory variants in K562 leukemia cells (Table S6).

Clearly, context-free WEVar has the overall best performance on the state-of-the-art independent testing datasets, which demonstrate its robustness and generality to predict functional noncoding variants across a wide range of context. Following WEVar, GWAVA_Unmatched, Unweighted average and FunSeq2 have superior performance to others. In contrast, CADD, DANN and FATHMM_MKL perform poorly. Particularly, FATHMM_MKL suffers from a low COR. Notably, integrating scores in a weighted scheme indeed boosts the prediction performance as demonstrated by the improvement of WEVar over Unweighted average.

### Context-dependent functional prediction

#### Overview of context-dependent WEVar

Different from context-free functional prediction, context-dependent functional prediction happens when a context-dependent WEVar is trained and the training and testing variants are from the same context. We develop “context-dependent” mode for WEVar because functional variants are usually studied in a cell type/tissue-specific way. The context-matching between training and testing variants may improve the prediction power. We demonstrate the prediction performance of context-free WEVar first, followed by a comparison between context-free and context-dependent WEVar to demonstrate the advantage for WEVar by being context-dependent.

#### Results of functional prediction between context-dependent WEVar and integrated scoring methods

We use the same benchmark datasets to evaluate context-free functional prediction. To restrict the training and testing variants from the same context, we randomly split each dataset into ten-folds with nine-folds as the training set and one-fold as the testing set. Tuning parameter for *L*_2_-norm is selected in the training set using 5-CV with AUROC as the evaluation metric. A final context-dependent WEVar is fitted using the whole training set with the selected tuning parameter and makes the functional prediction on the testing set. AUROC, AUPR and COR are calculated by comparing prediction scores and true labels of variants in the testing set. We use leave-one-fold-out by selecting nine-folds as training set and one-fold as testing set ten times. Accordingly, the whole procedure is repeated ten times and all evaluation metrics are reported as average.

We observe that context-dependent WEVar outperforms all scoring methods by obtaining the highest AUROC, AUPR and COR across all the benchmark datasets (Figure 4 and Table S1-S6). Moreover, we observe similar trends between context-dependent and context-free functional prediction, where WEVar, GWAVA_Unmatched and Unweighted average are the top-performed methods, while CADD, DANN and FATHMM_MKL have overall poor performance.

**Figure 4.**
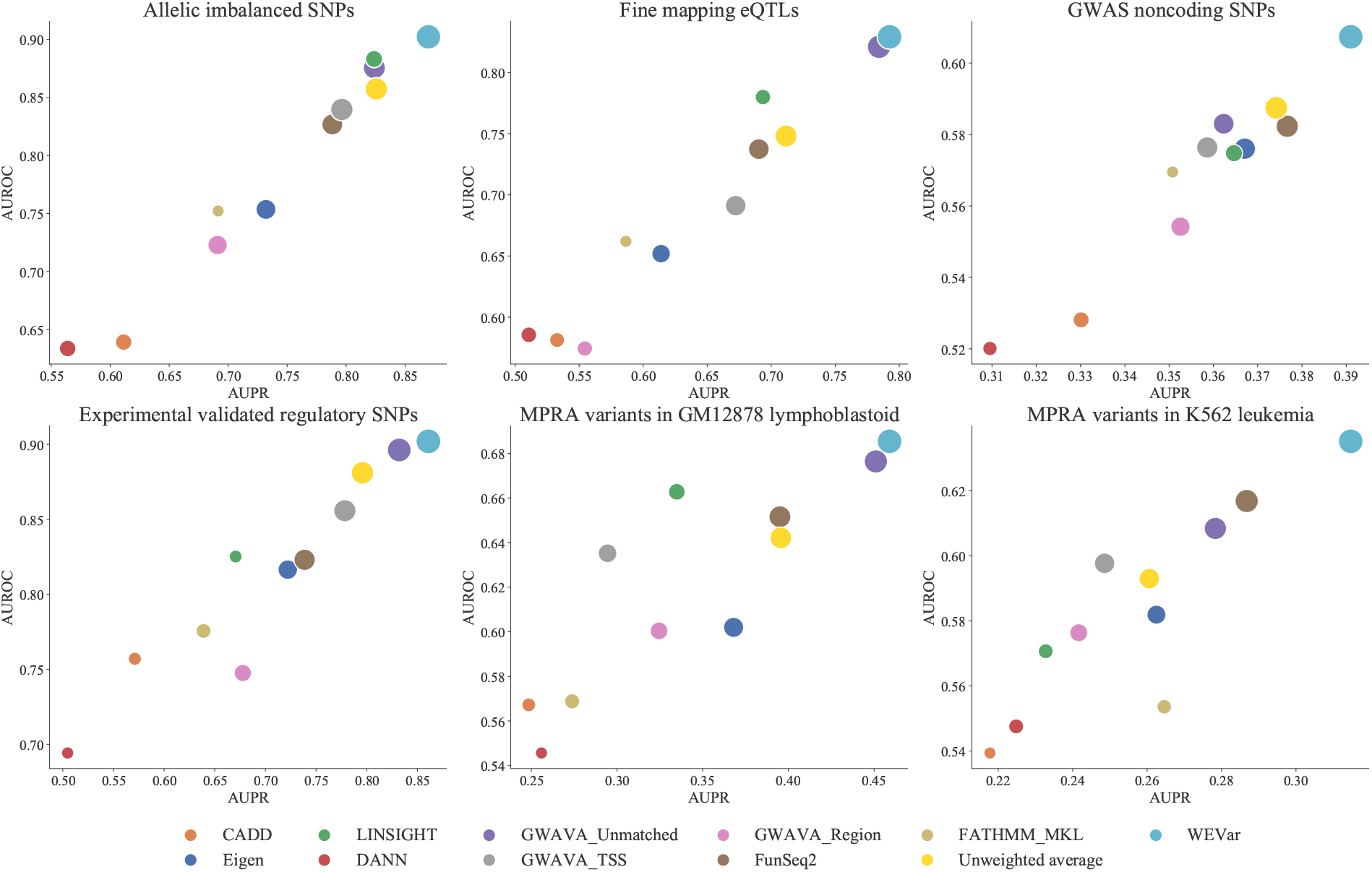
Evaluation of context-dependent WEVar and integrated scoring methods on state-of-the-art benchmark datasets, which include Allelic imbalanced SNPs, Fine mapping eQTLs, GWAS noncoding SNPs, Experimentally validated SNPs, MPRA validated variants in GM12878 lymphoblastoid cells and MPRA validated variants in K562 leukemia cells. We further remove variants on sex chromosome or with missing precomputed scores. To restrict the training and testing variants are from the same context, for each dataset, we randomly split the dataset into ten-folds with nine-folds as the training set and one-fold as the testing set. Context-dependent WEVar is trained on the nine-folds and independently evaluated on the left one-fold. AUC, AUCPR and COR are calculated and averaged in the ten replicates for each method. X axis presents AUPR; Y axis presents AUROC; the bubble size represents COR.

To further objectively gauge the performance of context-dependent WEVar, we utilize the training and testing variant set in the first part of challenge of Critical Assessment of Genome Interpretation eQTL challenge (CAGI) [34] derived from MPRA validated regulatory variants from GM12878 lymphoblastoid cells [32]. The variants selected by CAGI show significant level of transcriptional activity for either of two alleles. Specifically, the level of transcriptional activity is measured by differential abundance of transcripts versus plasmid input. Based on the FDR cutoff 0.01, a binary label is generated to indicate whether or not at least one of the two alleles of the variant exhibits a significantly high transcriptional activity (i.e. labeling 1 if FDR<0.01, otherwise, 0). As a result, the training set consists a total of 2,873 SNVs with 345 as positive set and 2,528 as negative set. The testing set contains a total of 2,808 SNVs with 348 positive variants and 2,460 negative variants. We further remove SNVs on sex chromosome or with missing precomputed scores in both sets. Besides following the original training and testing procedure, we further carry out an additional comparison by switching the training and testing set.

Consistent with our previous findings, context-dependent WEVar has superior performance to other scoring methods in both comparisons by achieving the highest AUROC, AUPR and COR, followed by GWAVA_Unmatched and Unweighted average (Figure 5, Table S7-8). Moreover, CADD and DANN have the overall poorest performance. The additional independent evaluation further strengthens the advantage of context-dependent WEVar in predicting functional noncoding variants by benefiting from matched context in training and testing set.

**Figure 5.**
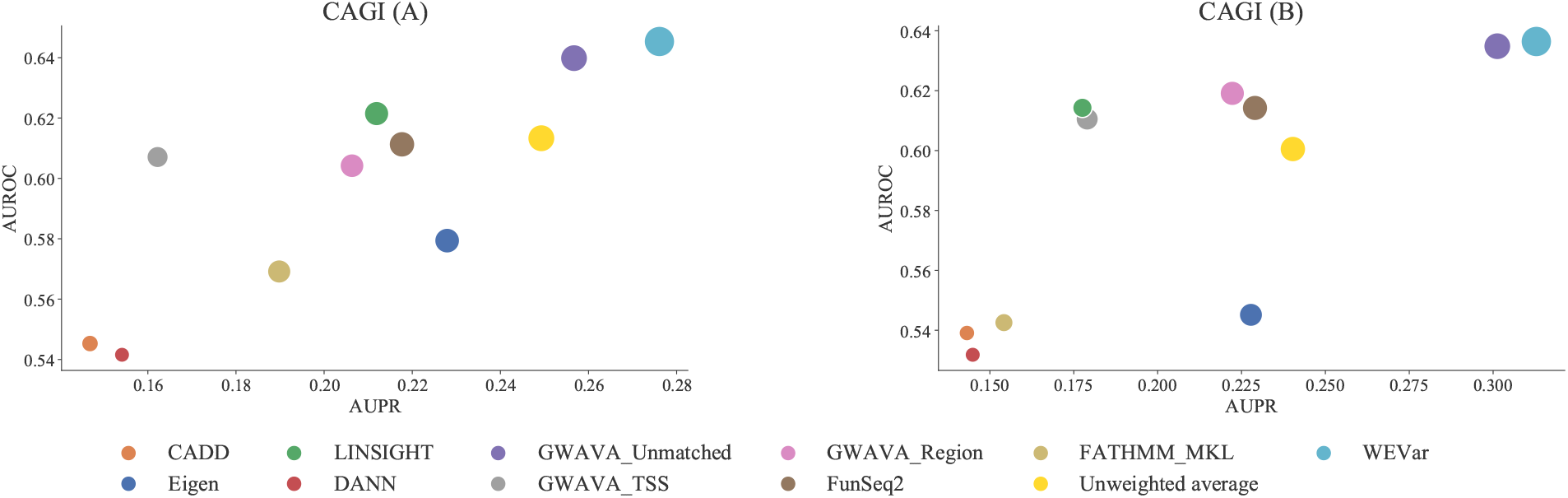
Prediction performance comparison between context-dependent WEVar and integrated scoring methods on the CAGI benchmark datasets. In CAGI, 2,873 SNVs with 345 as positive set and 2,528 as negative set. The testing set contains a total of 2,808 SNVs with 348 positive variants and 2,460 negative variants. We further remove SNVs on sex chromosome or with missing precomputed scores in both sets. (A) Context-dependent WEVar is first trained on the training set and evaluated on the testing set. (B) Similarly, we switch the training and testing set and perform an additional independent evaluation. The figure presents the AUPR, AUROC, and COR. X axis presents AUPR; Y axis presents AUROC; bubble size represents COR.

Besides improving the functional prediction, another important characteristic of WEVar is that it can identify the informative predictors that play the major contribution to the functional prediction among all integrated scoring methods. Consequently, we find that sets of informative predictors are different across benchmark datasets (Figure S1, Table S9). In most cases, WEVar identifies a parsimonious set of scoring methods that dominate the functional prediction especially FunSeq2 and GWAVA_Unmatched are two ubiquitous major contributors. Moreover, GWAVA_TSS is an additional major contributor for Allele imbalanced SNPs, Experimentally validated regulatory SNPs and integrated causal regulatory variants used by context-free WEVar. Regarding MPRA validated regulatory variants in GM12878 lymphoblastoid cells, Eigen is the additional method that has a major contribution. Similarly, GWAVA_Region and Eigen are two additional major contributors for two comparisons for CAGI training and testing variants. However, for GWAS noncoding SNPs and MPRA validated regulatory variants in K562 leukemia cells, there is a ubiquitous solution, where the contributions of all methods are relative uniform. These findings demonstrate that considering context-specificity in WEVar leads to different weight estimates and result in different sets of informative predictors. These observations also suggest that it is important to obtain an optimal weights when integrating different scoring methods, as the non-uniform weights estimated by WEVar lead improved functional prediction across benchmark datasets. Additionally, this point has been validated by both simulation and real data applications that WEVar outperforms the Unweighted average.

#### Results of comparison between context-free and context-dependent functional prediction

We hypothesize that considering context-specificity and context-matching context between training and testing variants in “context-dependent” WEVar will likely improve the predictive power for functional prediction. To validate this hypothesis, we directly compare the results of functional predictions between context-free and context-dependent WEVar on the aforementioned state-of-the-art benchmarking datasets (Figure S2, Table S1-S6).

For MPRA validated variants in GM12878 lymphoblastoid cells, context-dependent WEVar significantly outperforms context-free WEVar with large performance gain in around 5% AUPR and 8% COR but modest gain in AUROC. Similarly, context-dependent WEVar also achieves a large improvement by increasing about 4% AUPR and 4% COR but slightly improvement of AUROC for MPRA validated variants in K562 leukemia cells. Moreover, the improvement of context-dependent WEVar is evident demonstrated by nearly 5% and 3% increase in COR but slightly increase in AUROC and AUPR for both Fine mapping eQTLs and Allele imbalanced SNPs. In addition, context-dependent WEVar has a modest improvement of all metrics for GWAS noncoding SNPs. However, there is a lack of improvement on Experimentally validated regulatory SNPs, which could be explained by the small sample size of training set. This observation indicates that a large training set is necessary to improve the predictive power for context-dependent functional prediction. Overall, the comparisons between context-dependent and context-free WEVar validate the hypothesize that considering context-specificity and context-matching will improve the functional prediction. However, this improvement depends on the availability of enough sample size for training a robust context-dependent WEVar.

### Prioritization of causal regulatory variants by WEVar on benchmarking datasets

To demonstrate the application of WEVar in studying complex traits, we apply genome-wide functional scores of all noncoding variants in 1000 Genomes Project precomputed by context-free WEVar for finemapping analysis in risk loci. The diverse benchmarking datasets are generated from different experiments and study different traits, which are able to test the robustness of WEVar in prioritizing causal regulatory variants in risk loci.

#### Noncoding variants modulating gene expression

We evaluate WEVar on reported “expression-modulating variants” (emVars), which have been validated to show differential gene expression between alleles, from the MPRA study in GM12878 lymphoblastoid cells [32]. To assess whether these emVars with a strong linkage to GWAS SNPs can be prioritized by WEVar score, we create an extended LD block (*r*^2^ >0.2) utilizing ldproxy [35] to extract variants from all reference populations within the LD block, which are further assigned WEVar score.

Consequently, WEVar is able to prioritize emVars in exampled LD blocks (Figure S3 and Table S10). For example, emVar rs4790718 (chr17:4870893) scores higher than three LD-linked GWAS SNPs rs1060431 (chr17:4840868, pvalue=2×10^−26^), rs6065 (chr17:4836381, pvalue=2×10^−12^) and rs571461910 (chr17:4869143, pvalue=3.98 × 10^−9^), which are mapped to SPAG7 and associated with Platelet counts. Similarly, emVar rs922483 (chr8:11351912) is successfully prioritized by the highest score among all LD-linked variants including GWAS SNP rs2736340 (chr8:11343973, pvalue=6.03 × 10^−20^) associated with Systemic lupus erythematosus. Moreover, emVar rs56316188 (chr8:59323811) scores higher than GWAS SNP rs2859998 (chr8:59324162, pvalue=1 × 10^−7^), which is mapped to UBXN2B and associated with narcolepsy with cataplexy. Additionally, emVar rs306587 (chr10:30722908) is prioritized among LD-linked variants including one GWAS SNP rs1042058 (chr10:30728101, pvalue=6 × 10^−11^). Overall, these examples demonstrate that WEVar can successfully prioritize experimentally validated regulatory variants that modulate gene expression among LD-linked putatively causal GWAS SNPs, indicating that WEVar can potentially aid the fine mapping analysis.

#### Causal regulatory variants associated with Schizophrenia

Schizophrenia, typically diagnosed in the late teens years to early thirties, is a mental disorder characterized by disruptions in thought processes, perceptions, emotional responsiveness, and social interactions. Schizophrenia is one of the top 15 leading causes of disability worldwide [36, 37] and estimated international prevalence of schizophrenia among non-institutionalized persons is 0.33% to 0.75% [38]. Although GWAS has identified numerous noncoding schizophrenia-associated variants hypothesized to affect gene transcription, the causal regulatory variants are still elusive. To experimentally evaluate the regulatory potential of these GWAS SNPs and LD-linked variants, a recent study [39] screens several schizophrenia loci from a large GWAS cohort-Schizophrenia Working Group of the Psychiatric Genomics Consortium, using MPRA experiments in both K562 leukemia cells and SK-SY5Y neuroblastoma cells.

We apply context-free WEVar functional scores to discover causal regulatory variants associated with Schizophrenia. Briefly, we define “causal regulatory variants” as variants with significant differential expression between two alleles with a FDR cutoff 0.2. For each causal regulatory variant, we extend the risk locus by considering all variants in LD (*r*^2^ > 0.2). We further obtain precomputed context-free WEVar score for all variants in the risk locus. As a result, WEVar successfully prioritizes causal regulatory variants in the risk loci by assigning them the highest WEVar score (Figure 6 and Table S11). For example, rs34877519 (chr3:2554612) is successfully prioritized by obtaining the score higher than any variant in the risk locus including GWAS SNPs rs11708578 (chr3:2515894, pvalue=7.08 × 10^−11^) and rs17194490 (chr3:2547786, pvalue=1.00×10^−11^); rs7927437 (chr11:123395987) receives the highest score among all variants in the risk locus including GWAS SNP rs77502336 (chr11:123394636, pvalue= 3.98^−10^); rs7779548 (chr7:137074540) scores higher than any variant in the risk locus including GWAS SNP rs3735025 (chr7:137074844, pvalue=3.98× 10^−12^); rs6498914 (chr16:63699425) obtains the highest score among all variants in the risk locus including GWAS SNP rs2018916 (chr16:63700508, pvalue=7.08 × 10^−9^). Overall, these findings demonstrates that causal regulatory variants are not necessary the GWAS lead SNPs but the LD-linked variants. In addition, WEVar is a powerful tool in post-GWAS analysis to pinpoint the causal regulatory variants in the risk loci, which cannot be identified by a standard GWAS approach.

**Figure 6.**
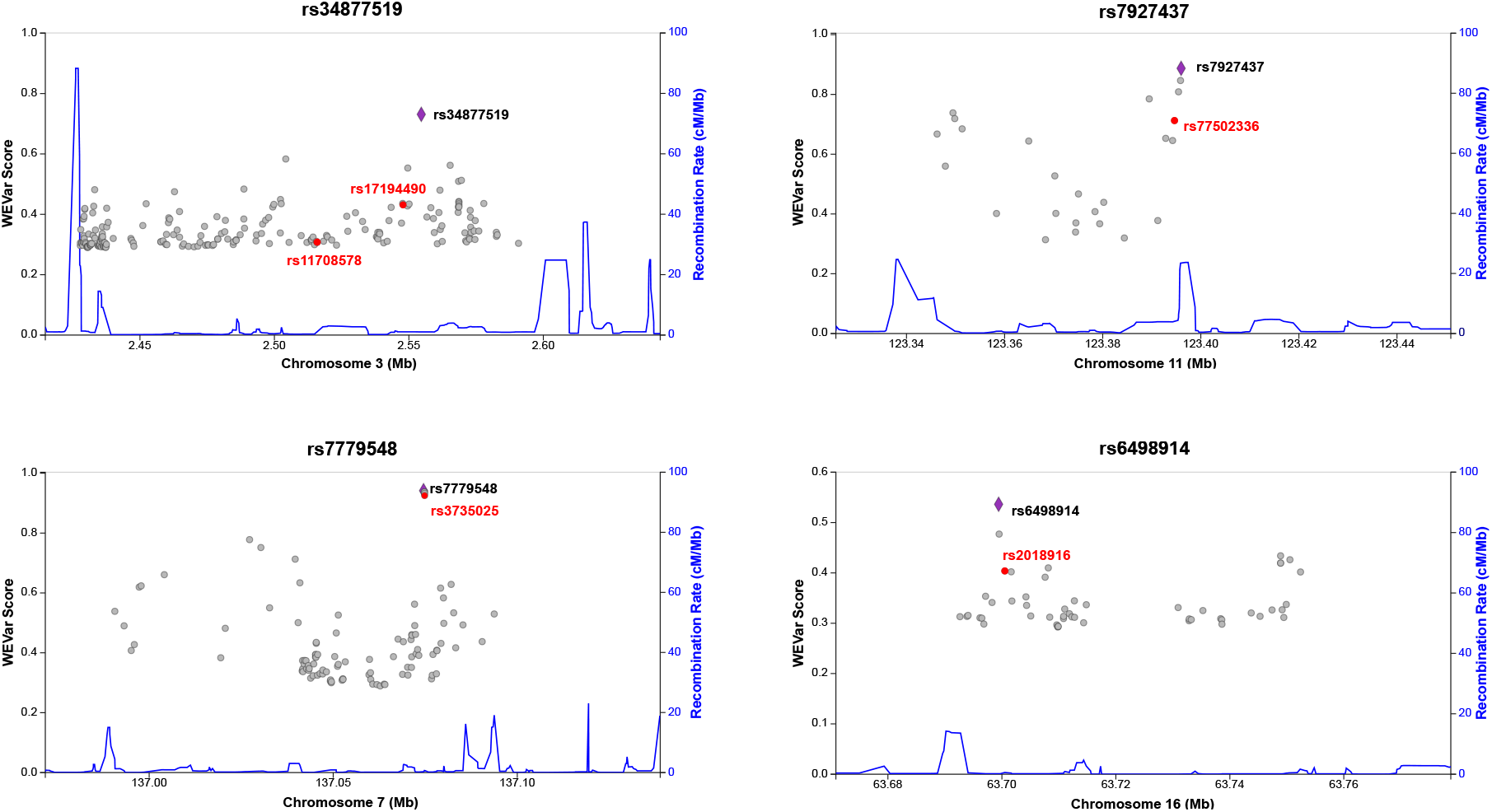
WEVar can prioritize causal regulatory variants associated with Schizophrenia. Causal regulatory variants are defined as variants with significant differential expression between two alleles (FDR<0.2) in MPRA experiments in both K562 leukemia cells and SK-SY5Y neuroblastoma cells. For each causal regulatory variant, we extend the risk locus by considering all variants in LD (*r*^2^ >0.2). As a result, rs34877519 (chr3:2554612) is successfully prioritized by obtaining the score higher than any variant in the risk locus including GWAS SNPs rs11708578 (chr3:2515894, pvalue=7.08 × 10^−11^) and rs17194490 (chr3:2547786, pvalue=1.00 × 10^−11^); rs7927437 (chr11:123395987) receives the highest score among all variants in the risk locus including GWAS SNP rs77502336 (chr11:123394636, pvalue=3.98 × 10^−10^); rs7779548 (chr7:137074540) scores higher than any variant in the risk locus including GWAS SNP rs3735025 (chr7:137074844, pvalue=3.98 × 10^−12^); rs6498914 (chr16:63699425) achieves the highest score among all variants in the risk locus including GWAS SNP rs2018916 (chr16:63700508, pvalue=7.08 × 10^−9^). The causal regulatory variants validated by MPRA are marked purple. LD-linked GWAS SNPs are marked red.

#### Causal regulatory variants associated with multiple traits and validated by multiple platforms

We benchmark WEVar on state-of-the-art datasets, which are generated from multiple studies for different traits such as Cleft lip/palate, heart, hair color and breast cancer and consists of regulatory variants experimentally validated by different functional assays. Similar to previous analyses, we define the risk locus by considering all variants in LD (*r*^2^ >0.2) for each regulatory variant. Consequently, WEVar is able to prioritize regulatory variants in each risk locus (Figure S4 and Table S12).

Specifically, rs6801957 (chr3:38767315, pvalue=9 × 10^−9^), located in intronic region of SCN10A, has been validated by BAC reporter system and 4C-seq to modulate cardiac SCN5A expression [40]. Consistent with the experimental validation, WEVar assigns the highest score to rs6801957 in the risk locus, which also includes multiple GWAS SNPs rs6795970 (chr3:38766675, pvalue=1 × 10^−58^), rs7433306 (chr3:38770639, pvalue=1× 10^−14^), rs6790396 (chr3:38771925, pvalue=2× 10^−39^), rs6599255 (chr3:38796415, pvalue=2×10^−^10) rs6798015 (chr3:38798836, pvalue=2×10^−12^) and rs10428132 (chr3:38777554, pvalue=1×10^−68^). We further evaluate another variant rs227727 (chr17:54776955, pvalue=7.3 × 10^−8^), which is mapped to 17q22 NOG locus and found associated with Cleft lip/palate. The NSCL/P-associated allele of rs227727 significantly decreases the nearby enhancer activity compared to the unassociated allele, which is experimentally validated by quantitative reporter assays transfected with a luciferase reporter vector [41]. Similarly, rs227727 is prioritized with the highest score in the risk locus. The next evaluated variant rs12821256 (chr12:89328335, pvalue=4×10^−30^) is located in a regulatory enhancer in the upstream of lncRNA LINC02458. It has been experimentally validated that rs12821256 is associated with hair color by altering the binding site of lymphoid enhancer-binding factor1 (LEF1) transcription factor. The altered binding site of LEF will reduce LEF responsiveness and enhancer activity in cultured human keratinocytes [42]. Again, rs12821256 scores highest in the risk locus, which is supported by the experimental finding. The last investigated variant is a breast cancer risk SNP rs11055880 (chr12:14410734), which resides in an intergenetic enhancer and validated by CRISPR-Cas9 approach [43] to have endogenous regulatory activities on expression of ATF7IP. Consistently, rs11055880 obtains the highest score among all variants in the risk locus.

Overall, the consistency between experimental validations and prioritization results based on WEVar score demonstrates the capability and robustness of WEVar to prioritize functional noncoding variants in a LD-linked risk locus. The robustness is reflected by the successful prioritization of heterogeneous variants, which are located in various genomic regions, associated with different traits, and validated by different functional assays.

## Discussion

In this work, we develop a statistical learning framework “WEVar” to predict functional noncoding variants by integrating representative scoring methods in an optimized weighted scheme. The development of WEVar is motivated by the existing gap of strong discordant performance of existing methods on state-of-the-art benchmark datasets, as shown by the inconsistent prediction performance of these methods on the integrated causal regulatory SNPs (Figure 2A).

Overall, the advantages of WEVar lies on several aspects. First, existing approaches, either supervised or unsupervised, are developed using thousands of functional annotations derived from multi-omics data deposited in large national consortia such as ENCODE and Roadmap Epigenomics. Different from existing methods, WEVar is developed on top of these methods by directly utilizing genome-wide precomputed functional scores, which collapse multi-dimensional functional annotations into a single score. Therefore, without losing information of functional annotations, direct application of the functional scores of existing approaches significantly reduces the dimensionality of feature space in model development of WEVar. Second, WEVar will identify informative predictors in an optimized weighted scheme and thus can leverage the advantages of different approaches, which likely lead to improved prediction performance compared to each integrated individual scoring method. Third, WEVar offers two modes: “context-free” and “context-dependent”. Each mode has its favorite scenario. We adopt a comprehensive training set [26], which integrates curated causal SNPs, located in different genomic regions, collected from different sources and associated with different traits to develop context-free WEVar. The large sample size, diverse context and genomic locations as well as heterogeneous trait association of these training variants make context-free WEVar powerful to predict functional noncoding variants with unknown or heterogeneous context. In contrast, training variant set of context-dependent WEVar is derived from the same context i.e. tissue-, cell type-, disease-specific. The context-specificity of training set makes context-dependent WEVar prefer the scenario when noncoding variants in training and testing set are from the same context, which may lead to improvement of functional prediction.

We perform a real data-based simulation study by considering the inherent correlations of precomputed functional scores among integrated scoring methods. The results demonstrate that WEVar outperforms individual scoring method and can estimate the contributions of integrated scoring methods accurately, which may explain the improved performance of WEVar. Next, we evaluate context-free functional prediction and context-dependent functional prediction respectively on state-of-the-art benchmark datasets, which include three variant sets containing putatively causal regulatory variants derived from statistical associations (i.e. Allelic imbalanced SNPs, Fine mapping eQTLs, GWAS significant noncoding SNPs), and three datasets consisting of experimentally validated regulatory variants (i.e. Experimentally validated regulatory SNPs, MPRA validated variants in GM12878 lymphoblastoid cells, MPRA validated variants in K562 leukemia cells). Besides evaluating context-dependent WEVar in each benchmark dataset by dividing it into training and testing set, we adapt an independent training and testing set from CAGI. Consequently, both context-free and context-dependent WEVar achieve an overall improvement of functional prediction compared to integrated scoring methods across all datasets. Specifically, WEVar outperforms Unweighted average, indicating the benefit of exploiting the optimized contributions of individual scoring method. GWAVA_Unmatched and Unweighted average are top-performed. In contrast, DANN, CADD and FATHMM_MKL always perform poorly. By comparing context-free and context-dependent WEVar on the same benchmark datasets, we find that context-dependent WEVar improve the functional prediction compared to context-free WEVar except for Experimentally validated regulatory SNPs possibly to the small sample size of training set. This observation indicates that being context-dependent improves the functional prediction and a large sample size is needed for make this improvement.

Another important characteristic of WEVar is that it can identify predictors that play major contribution to the functional prediction. As a result, major contributors are different across benchmark datasets. In most cases, WEVar identifies a parsimonious set of scoring methods that dominate the functional prediction especially FunSeq2 and GWAVA_Unmatched are two ubiquitous major contributors. However, for GWAS noncoding SNPs and MPRA validated regulatory variants in K562 leukemia cells, there is a ubiquitous solution, where the contributions of all methods are relative uniform. These findings demonstrate that both estimated weights and major contributors vary from context to context. Thus, it is important to obtain an optimal weights when integrating different scoring methods, as the non-uniform weights estimated by WEVar lead improved functional prediction across benchmark datasets. Additionally, this point is validated by both simulation and real data applications that WEVar outperforms the unweighted average of functional scores.

To demonstrate the application of WEVar in complex traits, we apply WEVar in the fine mapping analysis to evaluate whether it can successfully prioritize causal regulatory variants among LD-linked noncoding variants. By using precomputed WEVar score directly, variants assigned the highest score in a risk locus is considered to be prioritized. By using three benchmarking datasets of experimentally validated regulatory variants, we find that WEVar can prioritize regulatory variants modulating gene expression in GM12878 lymphoblastoid cells, associated with Schizophrenia and multiple traits such as Cleft lip/palate, heart, hair color and breast cancer. These findings demonstrate that WEVar can prioritize functional noncoding variants in risk loci and therefore alleviate the limitation of current GWAS, where the true causal SNPs may be masked by LD.

WEVar is a flexible approach, which can be further extended and improved by both integrated scoring methods and training variant set. In the current implementation, we include several representative scoring methods that are most popular in this field. With the rapid development post-GWAS analysis, there are other powerful methods developed or developing can be integrated into WEVar to further improve the prediction performance. The flexibility of WEVar is also reflected on the training variant set. With the affordability and popularity of functional assays such as massively parallel reporter assays (MPRAs) and clustered regularly interspaced short palindromic repeats (CRISPR)-based gene editing, more experimentally validated functional variants can be discovered and integrated into WEVar to improve the predictive power.

## Methods

WEVar is developed directly on top of precomputed functional score, which is an optimally integrative metric representing for thousands of functional annotations, from multiple individual scoring methods. Using these integrative functional scores directly will decrease the number of features in the model development and thus avoid the challenge high-dimensional data and multicollinearity. We will outline the details of WEVar in the following sessions.

### Obtaining precomputed functional scores

We download base-level genome-wide precomputed functional scores from all possible substitutions of single nucleotide variants (SNVs) in the human reference genome (GRCh37/hg19) from scoring methods including CADD [16], DANN [17], FunSeq2 [20], FATHMM-MKL [18], Eigen [21] and LINSIGHT [19]. In addition, we use three sets of precomputed scores from GWAVA (i.e. GWAVA_region, GWAVA_TSS, GWAVA_unmatched) for all SNVs in 1000 Genomes Project [22]. We choose these scoring methods to integrate into WEVar because they are widely used and mostly representative. Since the precomputed score of LINSIGHT is on region level, we assign all variants in the region with the same region-level score. More details for the source of these precomputed scores can be found in Table S13.

### Assembling variants in training and testing set

For context-free WEVar, the training variant set compiles a curated set of 5,247 causal regulatory variants including i) deleterious or pathogenic noncoding variants from the Human Gene Mutation Database (HGMD) [25] and ClinVar [44] ii) validated regulatory noncoding variants from the OregAnno [45] and candidate causal SNPs for 39 immune and non-immune diseases in the fine-mapping study [46] obtained from Li et al. [26]. The compiled variants are associated with different traits, have functional consequence in different tissues and cell types, and reside in different noncoding regions such as promoters, enhancers, 5’UTRs and 3’UTRs, making them ideal as a training set to predict functional consequence of noncoding variants from unknown or heterogeneous context. Accordingly, we collect six state-of-the-art benchmark independent variant sets from a wide range of context. Among them, three variant sets are collected from Li et al. [26], which include experimentally validated regulatory variants, expression quantitative trait loci (eQTL) [29] (FDR<0.1%) and allelic imbalanced SNPs [28] (FDR<0.1%) Moreover, GWAS significant noncoding SNPs are collected from NHGRI-EBI GWAS Catalog [30] (pvalue<10^−5^). Furthermore, two collected regulatory variant sets are validated by massively parallel reporter assays (MPRAs) in GM12878 lymphoblastoid cells [32] and K562 leukemia cells [33]. For context-free WEVar, these variant sets are used for independent testing. For context-dependent WEVar, we divide each variant set into ten folds with nine-folds as training set and one-fold as testing set.

### Statistical learning framework of WEVar

The workflow of WEVar is illustrated in Figure 1, which consists of four steps: (i) Creating the compiled training variant set (ii) Obtaining the precomputed functional scores for training variants (iii) Transforming the functional scores (iv) Training a constraint ensemble model.

#### Creating compiled training variant set

Depending on the purpose, we compile the training set for either “context-free” or “context-dependent” WEVar, as described in the section “Assembling variants in training and testing set”.

#### Obtaining precomputed functional scores for training variants

Precomputed functional scores are retrieved from representative scoring methods including CADD, DANN, Eigen, FunSeq2, FATHMM-MKL, LINSIGHT and GWAVA for variants in the training set, as described in the section “Obtaining precomputed functional scores”.

#### Transforming precomputed functional scores

Precomputed functional scores of integrated scoring methods are on different scales, which may result in different effect sizes of weight estimates by WEVar. However, the resulted different weight estimates are not due to different contributions of integrated scoring methods but because of the systematic bias induced by score scale. Therefore, it is important to perform a normalization step to make functional scores from different scales comparable. To integrate different scores are on the same scale, for each *j*th scoring method, we estimate two probability density functions (PDF) using kernel density estimation (KDE) based on the empirical distribution of the normalized scores for positive variant set and negative set respectively. As a result, PDF of the positive set denoted as **p**_*j*_(*s*|+) approximates the probability that a variant will have a prediction score *s* given the variant is functional (+), while PDF of the negative set denoted as **p**_*j*_(*s*|−) approximates the probability that a variant will have the same prediction score *s* given the variant is nonfunctional (−). Therefore, we use the ratio of two PDFs for the given *i*th variant, which is essentially the Bayes factor, to represent the likelihood the variant is functional versus nonfunctional. To stabilize the scale of the likelihood, we further take a logarithm of the ratio as the transformed score 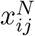 as:

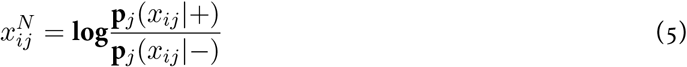

where *x_ij_* is the raw functional score of the *i*th variant in the *j*th scoring method; **p**_*j*_(*x_ij_*|+) and **p**_*j*_(*x_ij_*|−) are probability density of *x_ij_* in positive and negative set respectively.

#### Training a constraint ensemble model

Using the transformed scores, we will fit a constraint ensemble model, which is essentially a <u>C</u>onstrained <u>P</u>enalized <u>L</u>ogistic Regression model. Let **x**^*N*^ ∈ ℝ^*p*^ be the transformed scores of a variant for all scoring methods and *y* ∈ {−1, +1} be the variant label. The conditional probability of the variant being functional given **x**^*N*^ can be formulated as:

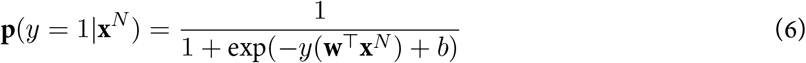

where **w**∈ ℝ^*p*^ is a weight vector, which contains the regression coefficients, and *b* ∈ *R* is the intercept. The likelihood function for *n* variants from both positive and negative set is defined as 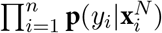. The objective function, which is the average of negative log-likelihood, is defined as:

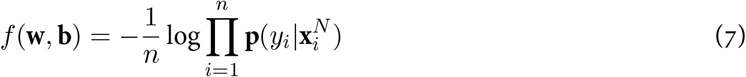

By minimizing the objective function, we can estimate **w** and *b* as:

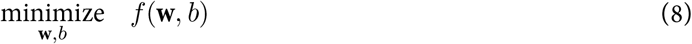

We further apply two constraints to the log-likelihood function. First, the weight of each scoring method is larger or equal to 0, indicating all scoring methods will contribute neutrally or positively to the prediction. Second, the sum of all weights equals to 1, which is a reasonable assumption for the summation of contributions from all scoring methods. In addition, to leverage all scoring methods by avoiding a sparse solution, we add an *L*_2_-norm to the objective function. Finally, we have the *L*_2_-norm regularized objective function with the two constraints as:

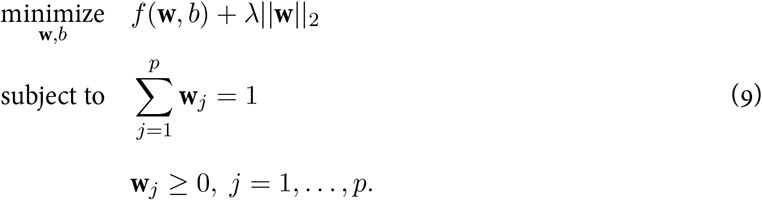

where *λ* ≥ 0 is the tuning parameter for *L*_2_-norm, which can be optimized from cross-validation in the training phase.

To minimize the loss function with equality and inequality constraints, we first rewrite the loss function as the standard form:

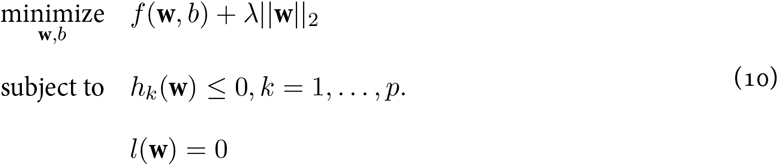

We then introduce Generalized Lagrange function to relax two constraints, which is formulated as:

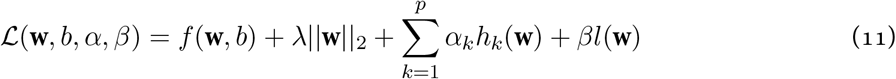

In this way, the dual problem is easier to solve compared with the primal problem. The primal solution can be constructed from the dual solution as:

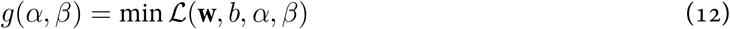

The Lagrange dual function can be considered as a pointwise maximization of some affine functions so it is always concave. The dual problem is always convex even if the primal problem is not convex, which can be easily solved by gradient-based methods.

#### Testing phase

In the testing phase, given variants be annotated precomputed functional scores from all scoring methods, which will be further transformed through the estimated KDE in the training phase. The transformed scores will serve as input features for trained ensemble model to predict the ensemble WEVar score.

#### Implementation

We adopt the SciPy [47], a Python scientific computing library, to perform the kernel density estimations, and CVXPY [48], a Python-embedded modeling library for convex optimization, to estimate constrained weights from the objective function.

#### Software availability

WEVar is implemented in a standalone software toolkit available at (https://github.com/lichen-lab/WEVar), which mainly consists of i) a compiled data package including precomputed scores for all SNVs (GRCh37/hg19) in 1000 Genomes Project across all integrated scoring methods; ii) a model package of pretrained context-free and context-dependent WEVar models; and iii) a Python software package to perform the functional prediction using pre-trained models or re-train a new model. To use a pre-trained model, WEVar will take compiled data package and genomic coordinates of testing variants as input. Alternatively, WEVar will take compiled data package and genomic coordinates of training variants to re-train a new WEVar model.

## Acknowledgments

This work was supported by the Indiana University Precision Health Initiative and Showalter Research Trust Fund [to L.C.] and the National Institute of Health R35 GM138342 [to Y.J.].

## Conflict of interest statement

None declared.

## Notes

### Competing Interest Statement

The authors have declared no competing interest.

